# Social investigation and social novelty in zebrafish: Roles of salience and novelty

**DOI:** 10.1101/2022.10.05.511022

**Authors:** Bianca Gomes do Nascimento, Caio Maximino

## Abstract

Social preference tests can be used to analyze variables that influence and modify social behaviors, and to investigate effects of substances such as medications, drugs, and hormones. They may become important tools for finding a valid model to study neuropsychiatric changes and to study human neurodevelopmental processes that have been impaired by social events. While a preference for conspecifics has been shown for different species, social novelty has been used as a model for anxiety-like behavior in rodents. The goal of this research was to understand the roles of stimulus salience (numerousness) and novelty in social investigation and social novelty tests in zebrafish (*Danio rerio* Hamilton 1822). We used a sequential design, in which animals are exposed first to a social investigation test (with dichotomous presentation of novel conspecifics vs. empty tank) and then to a social novelty test (with dichotomous presentation of the already known conspecific and a novel conspecific). In experiment 1, animals were presented to either 1 or 3 (vs. an empty tank) conspecifics as stimuli. In experiment 2, animals were presented to 1 vs. 3 conspecifics as stimuli. In experiment 3, animals were observed in the social investigation and social novelty tests for 3 consecutive days. The results showed equivalence between 1 or 3 conspecifics in the social investigation and social novelty tests, although animals were able to discriminate between different shoal sizes. These preferences do not change with repeated test exposure, suggesting novelty to be a minor contributor to social investigation and social novelty in zebrafish.

## 1 Introduction

In social animals, including zebrafish, social preference is a very common behavior, where the animal makes choices over their conspecifics (Liu et al., 2016). Social preference tests can be used to analyze variables that influence and modify social behaviors in vertebrates, and to investigate effects of substances such as drugs, toxicants, and hormones (Engeszer et al., 2004; Ogi et al., 2021). Furthermore, social preference tests can also be important tools to create valid models to study neuropsychiatric disorders and neurodevelopmental processes that impact social behavior (Maruska et al., 2019; Ogi et al., 2021; Soares et al., 2018).

The neuromolecular mechanisms underpinning social behavior are only likely to be fully understood using information obtained from various social, environmental, and physiological conditions (Oliveira, 2013; M. Taborsky et al., 2015). The first step in determining the structure of social behavior, thus, is to analyze preference and choices at the behavioral level.

Known as one of the most important model species in biology, the zebrafish is a small teleost belonging to the Cyprinidae family (Parichy, 2015) originating from tropical areas, mainly in India, and is becoming a strong name in areas of human diseases and in drug discovery and evaluation (Dodd et al., 2000; Gerlai, 2020; Stewart et al., 2014). Currently, the zebrafish is used as a model organism highly suitable for large-scale genetic manipulation, taking its importance to a higher level; furthermore, their embryos and larvae are even used for automated screening of high-yield drugs (Rinkwitz et al., 2011). With its numerous advantages regarding its use in laboratory research, the zebrafish is also an excellent tool for studies in the field of developmental biology thanks to its external development that allows visual analysis of the initiation performance of embryogenesis, enabling research with better results (Dooley & Zon, 2000; Roper & Tanguay, 2018).

The social behavior of zebrafish has been initially described from the point of view of shoaling, but soon the full complexity of sociality in this species was noticed (Maruska et al., 2019; Oliveira, 2013). Zebrafish are capable of cooperating with conspecifics (Pimentel et al., 2021) and of social learning (Lindeyer & Reader, 2010). The sight of conspecifics can be used as a reinforcer for zebrafish during operant conditioning (Al-Imari & Gerlai, 2008), suggesting that sociality is itself rewarding. Specific characteristics of conspecifics have been studied before, with zebrafish exhibiting social preferences for individuals similar in phenotype, age and size (Barba-Escobedo & Gould, 2012; Engeszer et al., 2004, 2007, 2008; Rosenthal & Ryan, 2005; Saverino & Gerlai, 2008). Social preferences in zebrafish are associated with many different neurotransmitter and neuromodulator systems, including dopamine (Scerbina et al., 2012), serotonin (Barba-Escobedo & Gould, 2012; Ponzoni et al., 2016), isotocin (Nunes et al., 2020; Ribeiro et al., 2020), and endocannabinoids (Barba-Escobedo & Gould, 2012).

Social preferences in zebrafish are visually mediated, with body morphology and body movement representing important dimensions that modulate preferences (Engeszer et al., 2008; Nunes et al., 2020). One of these dimensions is the salience of social stimuli. In a social preference test in which animals were presented with animated images of conspecifics, presentation of a single conspecific led to swimming away from the screen, while presenting up to 8 conspecifics led to a strong preference for swimming near the screen (Fernandes et al., 2015); however, no differences were observed between different stimulus numerosities (3, 5, or 7 conspecific images). Preferences for live conspecifics have been tested with different numerosities as well, with a preference for larger stimulus shoals (Potrich et al., 2015; Seguin & Gerlai, 2017). Baseline activity levels (body motion) seem to influence this response, as testing preference for conspecifics that are in warm water (high body motion) induce a marked preference for larger shoals, while testing preference for conspecifics that are in cold water (low body motion) abolishes this preference (Pritchard et al., 2001).

In the cases in which higher numerousness of social stimuli has been associated with social preferences, stimulus fish were also socially novel (Potrich et al., 2015; Pritchard et al., 2001; Seguin & Gerlai, 2017). However, Barba-Escobedo and Gould (2012) also showed that zebrafish display preference for social novelty, using a sequential design: in the first stage of the test (“social interaction”, SI), the choice presented is between a unfamiliar conspecific (obtained from a tank other than that of the focal fish) and an empty box, while in the second stage (“social novelty”, SN) the original conspecific (‘old stranger’) is presented in opposition to another unfamiliar conspecific (‘new stranger’); therefore, in the second stage the novelty of the ‘old stranger’ has already decreased, and the choice is between two conspecifics differing in novelty. Phenotype-dependent preferences for unfamiliar conspecifics were established, with AB and golden zebrafish showing preference both in the first stage (SI) and the second stage (SN)(Barba-Escobedo & Gould, 2012; Norton et al., 2019). The differences between social interaction and social novelty are relevant, given that zebrafish lacking a functional isotocin (oxytocin-like) receptor show normal preference in the SI test, but reduced preference in the SN test (Ribeiro et al., 2020). Moreover, behavior in the SN test is more correlated with novel object exploration than with social tendencies or behavior in the SI test across six zebrafish strains (Gonçalves et al., 2022). Gonçalves et al. (2022) suggested that preferences during the SN test are part of a “general exploration module”, while preferences in the SI test are more related to social tendencies.

The neurobehavioral differences in terms of social preference and novelty preference, as well as variations in responsiveness to social stimuli with differing salience (numerousness) point to the need to better understand the architecture of zebrafish social preferences in the SI and SN tests. The aim of the present study was to understand whether numerousness affects preference in both SI and SN tests, and whether decreasing novelty by repeated exposure to the conspecifics is sufficient to decrease preference in both tests.

## 2 Materials and methods

### 2.1 Animals and housing

82 adult zebrafish from the longfin phenotype were used in the experiments. Outbred populations were used due to their increased genetic variability, decreasing the effects of random genetic drift that could lead to the development of uniquely heritable traits (Parra et al., 2009; Speedie & Gerlai, 2008). The populations used in the present experiments are expected to better represent the natural populations in the wild. Animals were bought from a commercial vendor (Fernando Peixes, Belém/PA) and arrived in the laboratory with an approximate age of 3 months (standard length = 13.2 ± 1.4 mm), being quarantined for two weeks, before the experiment begin, when animals had an approximate age of 4 months (standard length = 23.0 ± 3.2 mm). Animals were kept in mixed-sex tanks during acclimation, with an approximate ratio of 1:1 males to females (confirmed by body morphology). The breeder was licensed for aquaculture under Ibama’s (Instituto Brasileiro do Meio Ambiente e dos Recursos Naturais Renováveis) Resolution 95/1993. Animals were group-housed in 40 L tanks, with a maximum density of 25 fish per tank, for the aforementioned quarantine before experiments began. Tanks were filled with non-chlorinated water at room temperature (28 °C) and a pH of 7.0-8.0. Lighting was provided by fluorescent lamps in a cycle of 14 hours of light-10 hours of darkness, according to standards of care for zebrafish (Lawrence, 2007).

Other water quality parameters were as follows: hardness 100-150 mg/L CaCO_3_; dissolved oxygen 7.5-8.0 mg/L; ammonia and nitrite < 0.001 ppm. Potential suffering of animals was minimized by controlling for the aforementioned environmental variables and scoring humane endpoints (clinical signs, behavioral changes, bacteriological status), following Brazilian legislation (Conselho Nacional de Controle de Experimentação Animal - CONCEA, 2017). Animals were used for only one experiment and in a single behavioral test, to reduce interference from apparatus exposure. Experiments were approved by UEPA’s IACUC under protocol 06/18.

### 2.2 Experiment 1: numerousness of social stimuli in preference and novelty experiments

32 animals were used in Experiment 1, with 24 being used as focal fish, and 8 being used as stimuli in the different stages and experimental conditions. To assess whether zebrafish are sensitive to the number of unfamiliar conspecifics in both SI and SN tests, fish were exposed to either one or three unfamiliar conspecifics in each stage. To perform this evaluation, in the social interaction phase, the experimental design consisted of 1 fish X no fish and 3 fish X no fish; in the social novelty phase, the experimental design consisted of 1 fish X 1 fish and 3 fish X 3 fish. A diagram of the experimental design can be found on Figure 1. Focal animals were transferred to glass tanks (13 × 13 × 17 cm) juxtaposed to two other glass tanks of equal dimensions, and visual access to these tanks was blocked by white plastic divisions. Animals were allowed to freely explore the tank for 5 min., after which the divisions were removed, allowing visual access to the side tanks. For half of the animals, one of the side tanks contained one unfamiliar conspecific, taken from a home tank different than that from which the focal animal was obtained. For the other half, one of the side tanks contained three unfamiliar conspecifics, again taken from a home tank different than that from which the focal animal was obtained. Since there were at least 10 manteinance tanks for any time during experiments, stimulus animals were drawn randomly from any of these tanks. Animals were allowed to freely explore the tank for another 10 minutes; this represents the *social interaction* (SI) stage. Immediately after that time period, the plastic divisions were inserted between tanks again, and the empty tank was quickly substituted by a tank containing the same amount of fish as the occupied one. Plastic divisions were then removed again, and the fish was allowed 10 more minutes to explore the tank; this represents the *social novelty* (SN) stage. In both the SI and SN stages, the side in which the “stimulus fish” was presented was randomly chosen between trials, and counter-balanced. Randomization was performed using the online tool at http://jerrydallal.com/random/assign.htm. Behavior was recorded with a video camera (Sony® DCR-DVD650 DVD Camcorder) mounted by the side of the tanks. Video files were then analyzed using automated tracking with TheRealFishTracker v. 0.4.0 (https://www.dgp.toronto.edu/~mccrae/projects/FishTracker/), running in Windows 10. While experimenters were not blinded to group, automated video tracking helps to reduce bias. The following variables were extracted for both SI and SN stages:

1. **Total time near stimulus fish/shoal** (SI) **or “new stranger”** (SN) (s), defined as the third of the focal tank nearest to the tank occupied by conspecifics. This is considered the primary endpoint of both tests.
2. **Total time near empty tank (**SI) **or “old stranger”** (SN) (s), defined as the third of the focal tank nearest to the unoccupied tank in the first stage, or the tank occupied by the previous unfamiliar fish in the second stage.
3. **Erratic swimming** (º), defined as the absolute turn angle averaged throughout the stage. This endpoint reflects an anxiety-like or fear-like component (Kalueff et al., 2013).
4. **Swimming speed** (cm/s), averaged throughout the stage.

**Figure 1.**
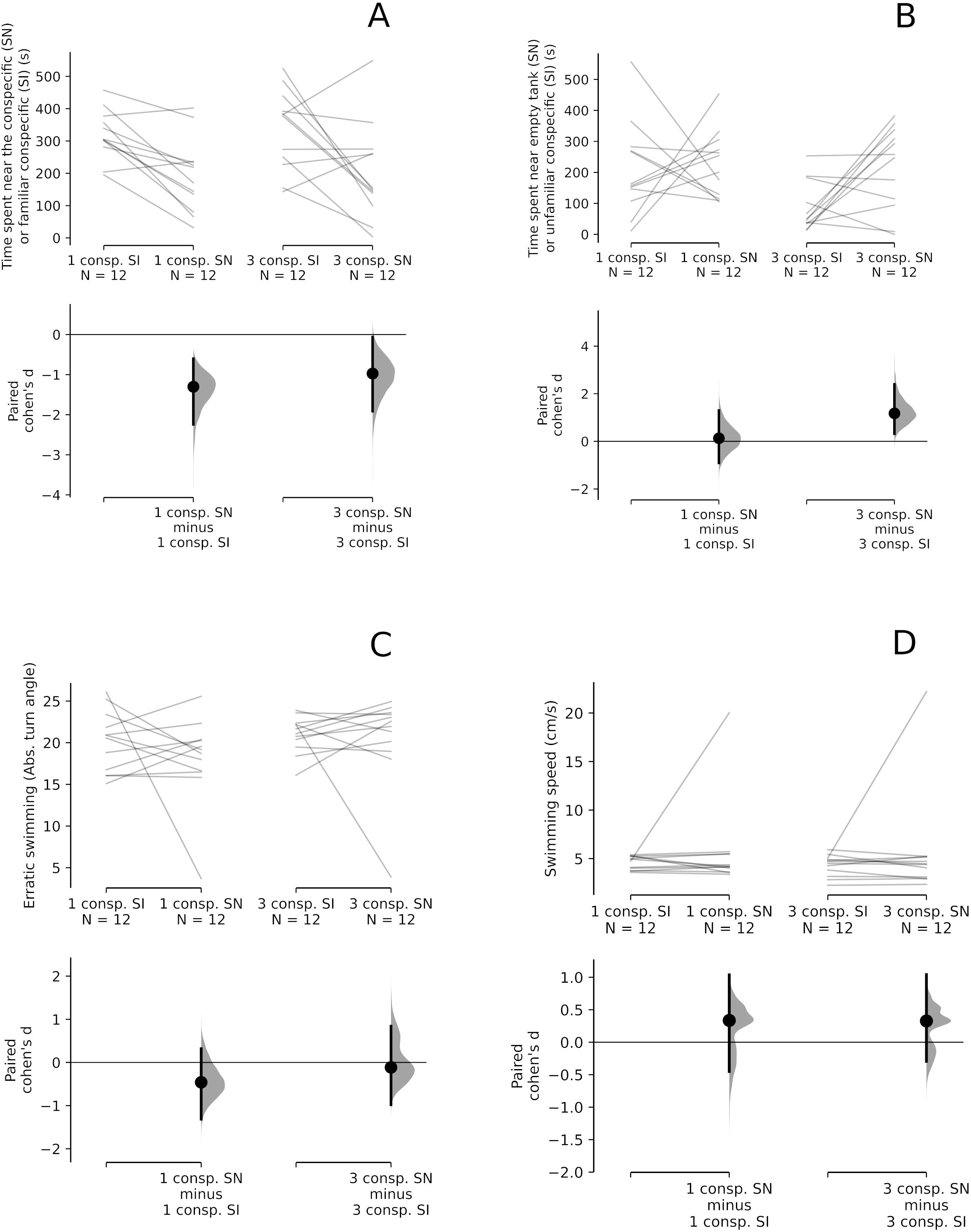
Experimental design

### 2.3 Statistical analysis for Experiment 1

To analyze differences between groups on the preference across SI and SN trials, repeated measures analyses of variance (ANOVAs) were applied, with test (SI vs. SN) as within-subjects variable and group as between-subjects variable. Since the assumption of normality was violated for most endpoints, we opted to use heteroscedastic one-way repeated measures ANOVA for trimmed means (Wilcox, 2012). *P*-values smaller than 0.05 were considered indicative of habituation. Effect sizes were calculated as partial eta-squared (η^2^p). Analyses were made in R 4.1.2, using the WRS2 package (v. 1.1-4, https://r-forge.r-project.org/projects/psychor/).

Data were represented using Cumming estimation plots (Ho et al., 2019), with individual datapoints plotted on the upper axes, with each paired set of observations (corresponding to the endpoint at either the SI or the SN test) connected by a line. On the lower axes, each paired mean difference was plotted as a bootstrap sampling distribution. Bootstrap resamples were taken using Efron’s (2000) method; the confidence interval is bias-corrected and accelerated. Plots were made using R 4.1.2, using the dabestr package (v. 0.3.0; https://github.com/ACCLAB/dabestr).

### 2.4. Experiment 2: Numerousness discrimination

The hypothesis of equivalence between groups in Experiment 1 could suggest that, in our experimental conditions, animals were not able to discriminate between conditions due to a lack of ability to discriminate quantities. Thus, in Experiment 2 we sought to understand whether that was the case. In order to do that, we exposed individual animals to the same apparatus and experimental design as in Experiment 1, except that groups differed in the choice that was available from the beginning of the experiment: one group were exposed to 1 fish X 1 fish, and another group was exposed to 3 fish X 1 fish. During the SN test, the fish in the side with smaller shoal was changed for another individual. A diagram of the experimental design can be found on Figure 1. Variables were analyzed as in Experiment 1, above. In Experiment 2, 32 animals were used in Experiment 1, with 24 being used as focal fish, and 8 being used as stimuli in the different stages and experimental conditions.

### 2.5 Statistical analysis for Experiment 2

To analyze differences between groups on the preference across SI and SN trials, repeated measures analyses of variance (ANOVAs) were applied, with test (SI vs. SN) as within-subjects variable and group as between-subjects variable. Since the assumption of normality was violated for most endpoints, we opted to use heteroscedastic one-way repeated measures ANOVA for trimmed means (Wilcox, 2012). *P*-values smaller than 0.05 were considered indicative of habituation. Effect sizes were calculated as partial eta-squared (η^2^p). Analyses were made in R 4.1.2, using the WRS2 package (v. 1.1-4, https://r-forge.r-project.org/projects/psychor/).

Data were represented using Cumming estimation plots (Ho et al., 2019), with individual datapoints plotted on the upper axes, with each paired set of observations (corresponding to the endpoint at either the SI or the SN test) connected by a line. On the lower axes, each paired mean difference was plotted as a bootstrap sampling distribution. Bootstrap resamples were taken using Efron’s (2000) method; the confidence interval is bias-corrected and accelerated. Plots were made using R 4.1.2, using the dabestr package (v. 0.3.0; https://github.com/ACCLAB/dabestr).

### 2.6 Experiment 3: Habituation of novelty

18 animals were used in Experiment 2, with 12 animals used as focal fish, and 6 as stimuli in the different conditions (SI and SN) and days. To understand the role of novelty in the SI and SN tests, animals were exposed to the tests again for 3 consecutive days, once a day, on all 3 testing days. This new exposure allowed us to evaluate the repeatability of the test. Animals were repeatedly exposed to the same procedure in Experiment 1, with a single conspecific as stimulus in both the SI and SN tests. A diagram of the experimental design can be found on Figure 1. Animals were isolated in individual 2 L tanks between days.

### 2.7 Statistical analysis for Experiment 3

To analyze the stability of preference and the role of novelty in SI and SN habituation, repeated measures analyses of variance (ANOVAs) were applied, with day and test as within-subjects variables. Since the assumption of normality was violated for most endpoints, we opted to use heteroscedastic one-way repeated measures ANOVA for trimmed means (Wilcox, 2012). *P*-values smaller than 0.05 were considered indicative of habituation. Effect sizes were calculated as δ_t_ (Algina et al., 2005). Analyses were made in R 4.1.2, using the WRS2 package (v. 1.1-4, https://r-forge.r-project.org/projects/psychor/).

## 3 Results

### 3.1 Experiment 1

No significant effect of group (1 fish vs. empty tank X 3 fish vs. empty tank) was found for the time spent near the conspecific (SI test) or familiar conspecific (SN test) (F(1, 22) = 0.12, p = 0.737, η^2^p = 0.01; Figure 2A). A significant within-subjects effect of test was found for this endpoint (F(1, 22) = 17.79, p < 0.001, η^2^p = 0.45), but no group X test interaction was found (F(1, 22) = 0.02, p = 0.898, η^2^p = 0.0); thus, for both groups, the time spent near the familiar conspecific in the SN trial was lower than the time spent near the conspecific in the SI trial, but no differences were found within trials.

**Figure 2.**
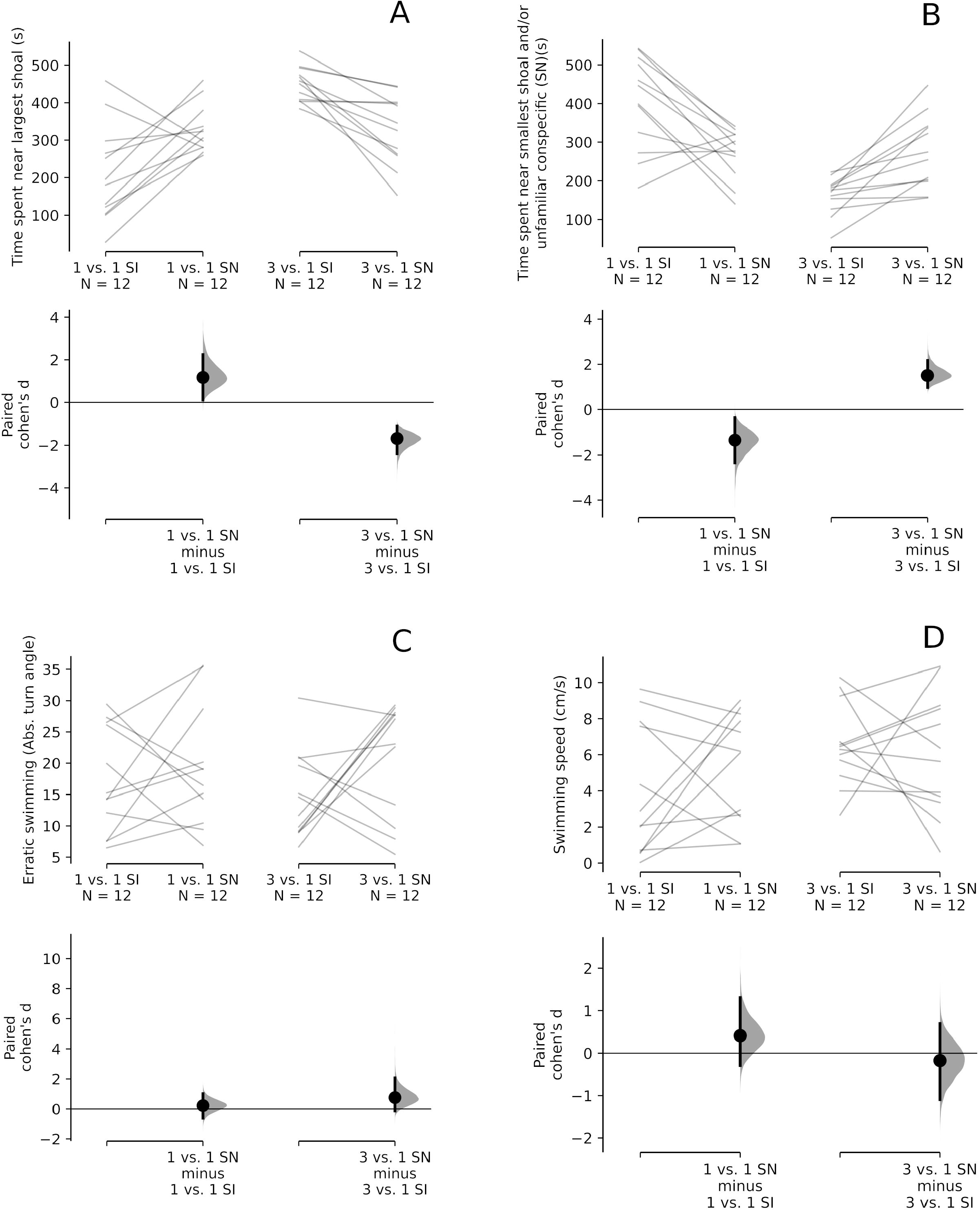
Lack of effect of salience/numerousness, indexed as number of conspecifics used as social stimuli, in the social investigation and social novelty tests. (A) Total time near stimulus fish/shoal (SI) or familiar conspecific (SN). (B) Total time near empty tank (SI) or unfamiliar conspecific (SN). (C) Erratic swimming. (D) Swimming speed. The effect sizes of differences between individuals exposed to 1 conspecific and 3 conspecifics vs. an empty tank in the social interaction and social novelty tests is shown in the Cumming estimation plots. The raw data is plotted on the upper axes; each paired set of observations is connected by a line. On the lower axes, each paired effect size (Cohen’s *d*) is plotted as a bootstrap sampling distribution. Mean differences are depicted as dots; 95% confidence intervals are indicated by the ends of the vertical error bars. 5000 bootstrap resamples were taken using Efron’s and Tibshinari’s (1994) method; the confidence interval is bias-corrected and accelerated.

Similarly, no main (between-subjects) effect of group was found for time near the empty tank (SI test) or unfamiliar conspecific (SN test) (F(1, 22) = 0.01, p = 0.936, η^2^p = 0.0; Figure 2B). A within-subjects effect of test (SI vs. SN) was found (F(1, 22) = 20.92, p < 0.001, η^2^p = 0.49), but a group X test interaction was absent (F(1, 22) = 0.2, p = 0.661, η^2^p = 0.01); thus, for both groups, the time spent near the unfamiliar conspecific in the SN trial was higher than the time spent near the empty tank in the SI trial, but no differences were found within trials.

No main (between-subjects) effect of group was found on erratic swimming (F(1, 22) = 2.08, p = 0.163, η^2^p = 0.09; Figure 2C). No within-subjects effect of test were found (F(1, 22) = 0.87, p = 0.362, η^2^p = 0.04), and no group X test interaction was found either (F(1, 22) = 0.33, p = 0.57, η^2^p = 0.01); thus, erratic swimming levels were similar both in the SI and SN trials for either group.

Finally, no main (between-subjects) effect of group was found on erratic swimming (F(1, 22) = 1.45, p = 0.241, η^2^p = 0.06; Figure 2D). No within-subjects effect of test were found (F(1, 22) = 0.03, p = 0.855, η^2^p = 0.00), and no group X test interaction was found either (F(1, 22) = 0.01, p = 0.935, η^2^p = 0.00); thus, swimming speeds were similar both in the SI and SN trials for either group.

### 3.2. Experiment 2

A main effect of group (1 conspecific vs. 1 conspecific X 3 conspecifics vs. 1 conspecific) was found for the time spent near the largest shoal (F(1, 22) = 19. 54, p < 0.001, η^2^p = 0.47; Figure 3A). No within-subjects test (SI vs. SN) effect was found (F(1, 22) = 0.01, p = 0.906, η^2^p = 0.00), but a group X test interaction was found (F(1, 22) = 24.9, p < 0.001, η^2^p = 0.53). Thus, in the 1 conspecific vs. 1 conspecific group, animals did not significantly spend time near any compartment during the SI test, but increased exploration of the tank with the unfamiliar conspecific in the SN test; while in the 3 conspecific vs. 1 conspecific group, animals spend more time near the largest shoal during the SI test, but decreased exploration of the tank with the familiar conspecifics (i.e., the largest shoal) in the SN test.

**Figure 3.**
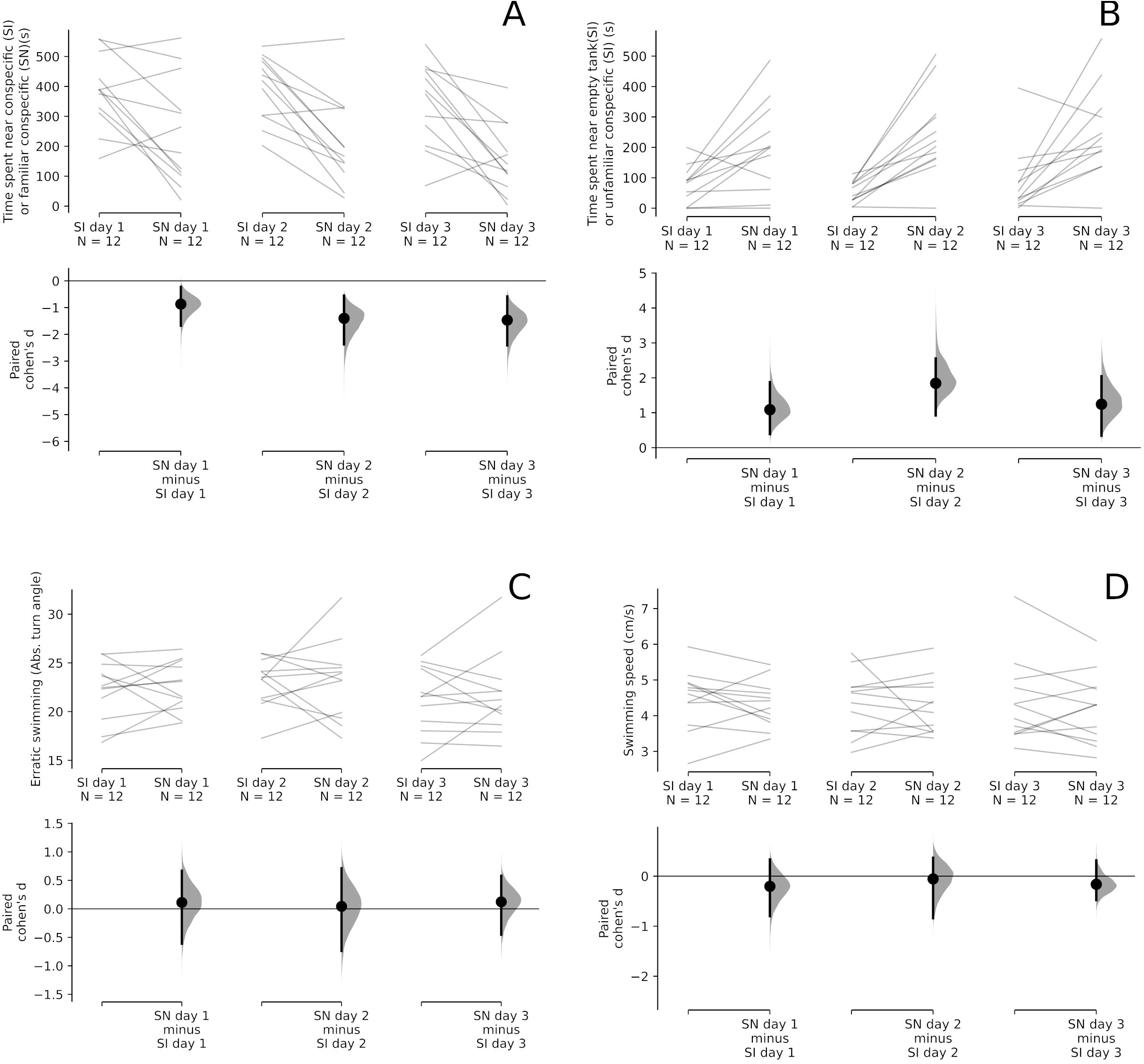
Discrimination of numerousness in the social investigation and social discrimination tests in animals exposed to different conspecific numbers in both tanks. (A) Total time near largest stimulus fish (shoal). (B) Total time near smallest stimulus (single fish). (C) Erratic swimming. (D) Swimming speed. The effect sizes of differences between individuals exposed to 1 conspecific vs. 3 conspecifics in the SI and SN tests is shown in the Cumming estimation plots. The raw data is plotted on the upper axes; each paired set of observations is connected by a line. On the lower axes, each paired effect size (Cohen’s *d*) is plotted as a bootstrap sampling distribution. Mean differences are depicted as dots; 95% confidence intervals are indicated by the ends of the vertical error bars. 5000 bootstrap resamples were taken using Efron’s and Tibshinari’s (1994) method; the confidence interval is bias-corrected and accelerated.

Similarly, a main effect of group (1 conspecific vs. 1 conspecific X 3 conspecifics vs. 1 conspecific) was found for the time spent near the smallest shoal (i.e., 1 conspecific) (F(1, 22) = 19. 72, p < 0.001, η^2^p = 0.47; Figure 3B). No within-subjects test (SI vs. SN) effect was found (F(1, 22) = 0.17, p = 0.683, η^2^p = 0.01), but a group X test interaction was found (F(1, 22) = 27.33, p < 0.001, η^2^p = 0.55). Thus, in the 1 conspecific vs. 1 conspecific group, animals did not significantly spend time near any compartment during the SI test, but decreased exploration of the tank with the familiar conspecific in the SN test; while in the 3 conspecific vs. 1 conspecific group, animals spend less time near the smallest shoal during the SI test, but increased exploration of the tank with the unfamiliar conspecifics (i.e., the smallest shoal) in the SN test.

No main (between-subjects) effect of group was found on erratic swimming (F(1, 22) = 0.03, p = 0.863, η^2^p = 0.00; Figure 3C). No within-subjects effect of test were found (F(1, 22) = 2.52, p = 0.127, η^2^p = 0.1), and no group X test interaction was found either (F(1, 22) = 0.66, p = 0.426, η^2^p = 0.03); thus, erratic swimming levels were similar both in the SI and SN trials for either group.

Finally, no main (between-subjects) effect of group was found on erratic swimming (F(1, 22) = 3.29, p = 0.083, η^2^p = 0.13; Figure 3D). No within-subjects effect of test were found (F(1, 22) = 0.25, p = 0.623, η^2^p = 0.01), and no group X test interaction was found either (F(1, 22) = 1.14, p = 0.297, η^2^p = 0.05); thus, swimming speeds were similar both in the SI and SN trials for either group.

### 3.3 Experiment 3

A main within-subjects effect of test (SI vs. SN) was found for the time spent near the conspecific (SI test) or the familiar conspecific (SN test)(F(1, 11) = 81.94, p < 0.001, η^2^p = 0.88; Figure 4A). No within-subjects effect of day was found for this variable (F(2, 22) = 1.35, p = 0.279, η^2^p = 0.11), and a test X day interaction was also absent (F(2, 22) = 0.4, p = 0.677, η^2^p = 0.03). Thus, animals spent more time near the conspecific than the empty tank in the SI test and far from the familiar conspecific in the SN test regardless of test day.

**Figure 4.**
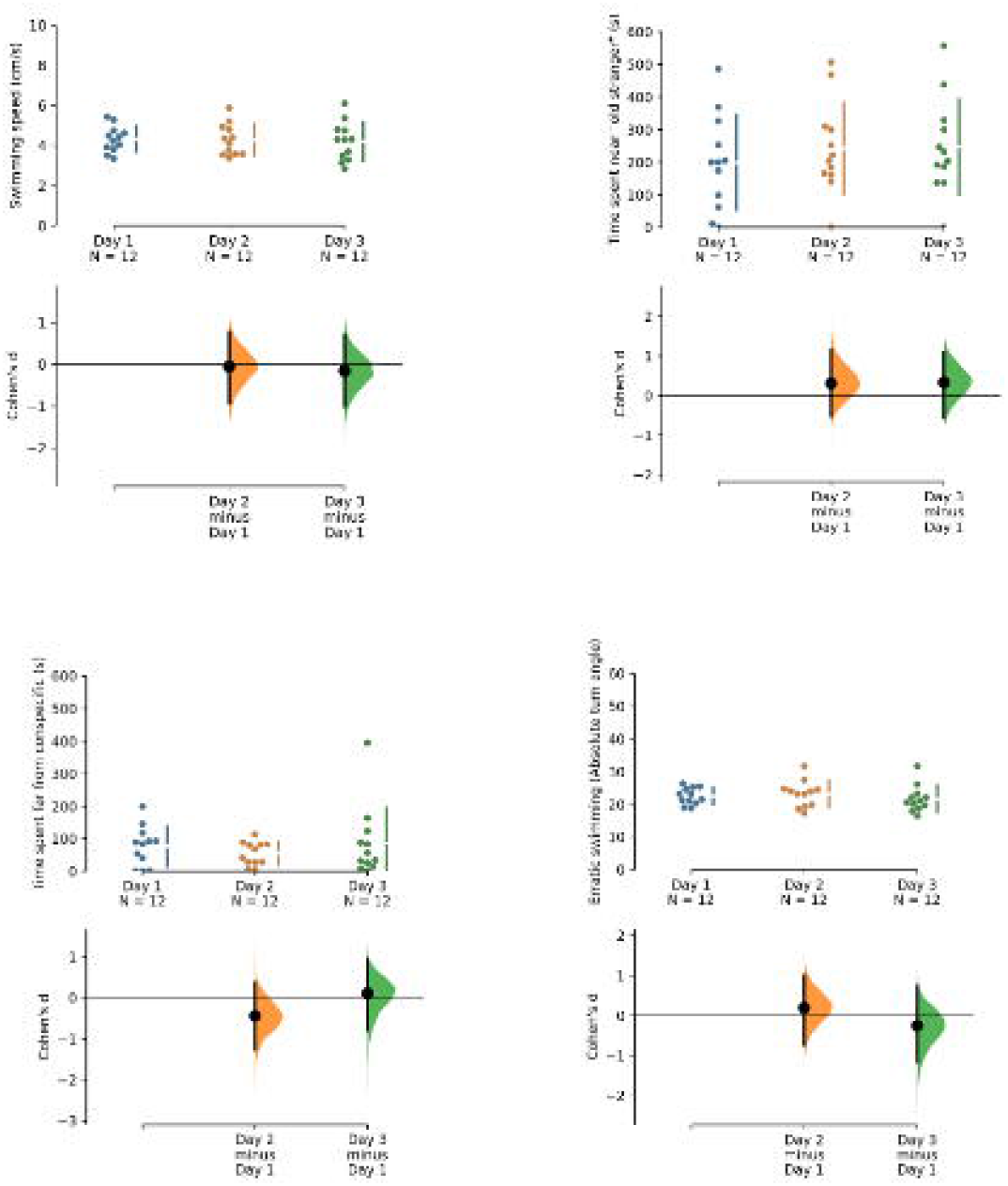
Stability of social preference in the social investigation and social novelty tests after repeated exposure for three days. (A) Total time near stimulus fish (SI) or familiar conspecific (SN). (B) Total time near empty tank (SI) or unfamiliar conspecific (SN). (C) Erratic swimming. (D) Swimming speed. The raw data is plotted on the upper axes; eachpaired set of observations (SI vs. SN) is connected by a line. On the lower axes, each paired mean difference is plotted as a bootstrap sampling distribution. Mean differences are depicted as dots; 95% confidence intervals are indicated by the ends of the vertical error bars. Each mean difference is depicted as a dot. Each 95% confidence interval is indicated by the ends of the vertical error bars. 5000 bootstrap resamples were taken using Efron’s and Tibshinari’s (1994) method; the confidence interval is bias-corrected and accelerated.

Similarly, a within-subjects effect of test (SI vs. SN) was found for time near the empty tank (SI test) or near the unfamiliar conspecific (SN test)(F(1, 11) = 53.94, p < 0.001, η^2^p = 0.83; Figure 4B). No within-subjects effect of day was found for these variables (F(2, 22) = 0.31, p = 0.737, η^2^p = 0.03), and a test X day interaction was also absent (F(2, 22) = 0.61, p = 0.555, η^2^p = 0.05). Thus, animals spent less time near the empty tank in the SI test and more time near the unfamiliar conspecific in the SN test, and these endpoints showed no habituation.

No within-subjects effect of test (SI vs. SN) was found for erratic swimming (F(1, 11) = 0.49, p = 0.5, η^2^p = 0.04; Figure 4C). Similarly, no within-subjects effect of day was found (F(2, 22) = 1.43, p = 0.261, η^2^p = 0.12), and no test X day interaction was found as well (F(2, 22) = 0.02, p = 0.978, η^2^p = 0.00).

Finally, no within-subjects effect of test (SI vs. SN) was found for swimming speed (F(1, 11) = 1.68, p = 0.221, η^2^p = 0.13; Figure 4D). Similarly, no within-subjects effect of day was found (F(2, 22) = 0.16, p = 0.856, η^2^p = 0.01), and no test X day interaction was found as well (F(2, 22) = 0.1, p = 0.909, η^2^p = 0.01).

## 4 Discussion

The present experiments assessed the role of numerousness and habituation in the zebrafish social interaction (SI) and social novelty (SN) tests. It was found that zebrafish do not show stronger preferences for 3 conspecifics vs. an empty tank than 1 conspecific vs. an empty box in neither the SI test nor the SN test. Nonetheless, evidence for preference for conspecifics was observed in all experiments, and these preferences did not change with repeated exposure to the test apparatuses.

Preference for larger numerosities has been shown in other studies in fish, usually as evidence of the cognitive capacities of teleosts (Potrich et al., 2015; Seguin & Gerlai, 2017). In the present experiments, we were interested in whether preferences were stronger for individuals presented with a single conspecific or a shoal, as an index of social stimulus salience. Thus, we were interested in whether zebrafish would adjust their social preferences depending on the numerousness of the stimulus. In the SI test – in which the individual chooses between a social stimulus and an empty tank (Barba-Escobedo & Gould, 2012) – preferences for either 1 or 3 conspecifics was equivalent (Experiment 1). However, when animals were exposed to 1 or 3 conspecifics simultaneously (Experiment 2), a preference for the larger shoal was found. Thus, both Experiments 1 and 2 strengthen the observation that zebrafish are strongly motivated to investigate conspecifics, irrespective of shoal sizes. Social stimulus novelty is an important factor in this test, however, and the presence of a larger shoal could represent not a more reinforcing stimulus, but simply a more novel social context (since the number of novel stimuli to investigate would be higher). In the SN test, individuals must choose between the conspecific which was previously shown and a novel conspecific; thus, the novelty of the previous interaction decreases, while individuals are shown another social stimulus with higher novelty. In that situation, the novelty of the first unknown conspecific (“stranger 1”) decreases, while the novelty of the second unknown conspecific (“stranger 2”) is at its highest.

Another possible source of variability in our experimental designs is that, across all trials and conditions, stimulus fish were able to establish visual contact with the focal fish. In cases in which a single stimulus fish was presented, it is likely that this individual responded to the behavior of the focal fish by either approaching the fish or presenting agonistic/aggressive displays; in its turn, the focal fish could respond to the stimulus fish’s behavior. Thus, the responsiveness of the stimulus fish, and not only novelty or salience, could be important in establishing the preferences in both the SI and SN stages of Experiment 1. Conversely, this could not occur in the 3-fish treatment because the 3 stimuli interacted mostly between themselves. We have not quantified the behavior of stimulus fish, given that this would necessitate other more robust methods. However, this remains a possibility, given that Pritchard et al. (2001) showed that the levels of body motion of stimulus fish influence social preference in zebrafish. Since no differences were found between groups in Experiment 1, however, it is possible that this behavioral interaction was not the main driver of preferences in the present experiments.

We found that increasing stimulus salience did not lead to higher preference during both stages (SI and SN). Two interpretations are equally possible: that, in situations with larger numerosities, novelty discrimination is harder; or that social novelty is a stronger motivator for social preference than salience in zebrafish. Experiment 2 discarded the possibility that, under the conditions of the present experiments, animals were simply not able to discriminate between different quantities, as animals preferred the larger shoal vs. 1 individual in both the SI and SN phases. Nonetheless, how familiarity is established and how long it takes for zebrafish to establish discriminate between unfamiliar/novel and familiar conspecifics is still unknown. As discussed below, our results suggest that social recognition can be established as quickly as 10 min., and a social recognition memory is maintained for at least 24 h. Animals were kept in maintenance tanks for at least two weeks before experiments begun, an interval that is sufficient to establish familiarity in guppies (Griffiths & Magurran, 1997). Thus, animals in the present experiments could use cues from this previous experience to discriminate between familiar and unfamiliar individuals, but the possibility that this discrimination is harder for larger shoals (larger numerosities).

Novelty is also anxiogenic for most vertebrates (Maximino et al., 2012; Montgomery, 1955; Montgomery & Monkman, 1955); novel social situations are inherently anxiogenic, since they involve an approach-avoidance conflict in which potential benefits of social interactions with novel individuals are counterbalanced by potential risks (Soares et al., 2018). Socially competent individuals of a given species are more likely to be sensitive to the risk-benefit balance and assess the situation, plastically changing its behavior to better adapt the actual social context (B. Taborsky & Oliveira, 2012). Nonetheless, preference was found to be equivalent after repeated exposures to the apparatus (that is, preference in the SI test did not habituate) or to social novelty (that is, preference in the SN test also did not habituate). Moreover, no effects were observed of either salience/numerousness (Experiment 1) or habituation (Experiment 3) on typical measures of anxiety-like behavior, such as erratic swimming and freezing.

While the role of novelty has been investigated previously mainly in relation to the SN test, social novelty also seems to play a role in SI as well. Norton et al. (2019) used a similar design (with a sequential SI-SN design) to test preferences in zebrafish from the AB line. While results from the SN were similar to ours and to Barba-Escobedo’s and Gould’s (2012), with animals preferring to investigate unfamiliar conspecifics than those presented during the SI phase; however, animals also showed a stronger preference for unfamiliar conspecifics over an empty tank during the SI phase (Norton et al., 2019). The importance of social novelty in social investigation is inconsistent with findings from our Experiment 3, however, as decreasing novelty did not decrease preference. A possible explanation is differences in lineage used; while Norton et al. (2019) used fish from the AB lineage, we used wild-type longfin animals. Differences in sociality between AB and other outbred animals (e.g., wild-type shortfin zebrafish) were observed by Barba-Escobedo and Gould (2012), with AB zebrafish showing a strong preference for an unfamiliar conspecific in both the SI and SN tests, while shortfin zebrafish showed a preference for the empty tank in the SI test and no preference in the SN test. Scerbina et al. (2012) showed that AB zebrafish also show a stronger preference for computer animated images in a SI test than shortfin zebrafish. Thus, it is possible that AB zebrafish are more social and/or more sensitive to social novelty than outbred populations. Unfortunately, a direct comparison between lineages was not possible in our case.

The lack of habituation across three trials separated by 24 h also suggest that zebrafish are able to retain the memory of the conspecific for at least 24 h. Indeed, Madeira and Oliveira (2017) showed that animals from the Tu wild-type strain spend more time exploring an unfamiliar conspecific 24 h after being exposed to a SI test (with 1 unfamiliar conspecific in each compartment). This suggests that the information about familiarity/novelty was retained for at least 24 h, similar to the present experiments. Experiments with longer exposures and/or intervals can elucidate how long this social recognition memory lasts in zebrafish.

Taken together, these results suggest that social preference is not guided by stimulus salience and novelty in zebrafish. These results underline the complexity of zebrafish social behavior and the need to understand the motivational basis of this process. Moreover, it also points to practical considerations on using social preference (interaction and novelty) tests in neurobehavioral research. For instance, since social preferences do not appear to be changed by numerousness when individuals need to choose between conspecifics and an empty tank, it is possible to reduce the number of animals used in the experiments. Moreover, since preference does not seem to habituate, repeated exposure can be made using the same individuals, which is advantageous to assess the effects of repeated treatments (including those that are common in pharmacological research). The separability between numerousness discrimination and social preference also suggests avenues for investigating the brain mechanisms relating to both capabilities, including regions related to cognitive vs. motivational functions.

## References

Al-Imari, L., & Gerlai, R. (2008). Sight of conspecifics as reward in associative learning in zebrafish (Danio rerio). Behavioural Brain Research, 189(1), 216–219. https://doi.org/10.1016/j.bbr.2007.12.007

Barba-Escobedo, P. A., & Gould, G. G. (2012). Visual social preferences of lone zebrafish in a novel environment: Strain and anxiolytic effects. Genes, Brain and Behavior, 11(3), 366–373. https://doi.org/10.1111/j.1601-183X.2012.00770.x

Cohen, J. (1988). Statistical power analysis for the behavioral sciences (2nd ed). Routledge.

Conselho Nacional de Controle de Experimentação Animal - CONCEA (2017). Diretriz brasileira para o cuidado e a utilização de animais para fins científicos e didáticos—DBCA. Anexo I. Peixes mantidos em instalações de instituições de ensino ou pesquisa científica, Resolução Normativa CONCEA no 34 218 (2017). https://www.mctic.gov.br/mctic/opencms/legislacao/outros_atos/resolucoes/Resolucao_Normativa_CONCEA_n_34_de_27072017.html

Dodd, A., Curtis, P. M., Williams, L. C., & Love, D. R. (2000). Zebrafish: Bridging the gap between development and disease. Human Molecular Genetics, 9(16), 2443–2449. https://doi.org/10.1093/hmg/9.16.2443

Dooley, K., & Zon, L. I. (2000). Zebrafish: A model system for the study of human disease. Current Opinion in Genetics & Development, 10(3), 252–256. https://doi.org/10.1016/S0959-437X(00)00074-5

Efron, B. & Tibshirani, R. J. (1994) An Introduction to the Bootstrap. Boca Raton: CRC Press.

Engeszer, R. E., Da Barbiano, L. A., Ryan, M. J., & Parichy, D. M. (2007). Timing and plasticity of shoaling behaviour in the zebrafish, Danio rerio. Animal Behaviour, 74(5), 1269–1275. https://doi.org/10.1016/j.anbehav.2007.01.032

Engeszer, R. E., Ryan, M. J., & Parichy, D. M. (2004). Learned Social Preference in Zebrafish. Current Biology, 14(10), 881–884. https://doi.org/10.1016/j.cub.2004.04.042

Engeszer, R. E., Wang, G., Ryan, M. J., & Parichy, D. M. (2008). Sex-specific perceptual spaces for a vertebrate basal social aggregative behavior. Proceedings of the National Academy of Sciences, 105(3), 929–933. https://doi.org/10.1073/pnas.0708778105

Fernandes, Y., Rampersad, M., Jia, J., & Gerlai, R. (2015). The effect of the number and size of animated conspecific images on shoaling responses of zebrafish. Pharmacology Biochemistry and Behavior, 139, 94–102. https://doi.org/10.1016/j.pbb.2015.01.011

Gerlai, R. (2020). Evolutionary conservation, translational relevance and cognitive function: The future of zebrafish in behavioral neuroscience. Neuroscience & Biobehavioral Reviews, 116, 426–435. https://doi.org/10.1016/j.neubiorev.2020.07.009

Gonçalves, C., Kareklas, K., Teles, M. C., Varela, S. A. M., Costa, J., Leite, R. B., Paixão, T., & Oliveira, R. F. (2022). Phenotypic architecture of sociality and its associated genetic polymorphisms in zebrafish. Genes, Brain and Behavior, 21(5), e12809. https://doi.org/10.1111/gbb.12809

Griffiths, S. W. & Magurran, A. E. (1997). Familiarity in schooling fish: How long does it take to acquire? Animal Behaviour, 53, 945–949. https://doi.org/10.1006/anbe.1996.0315

Ho, J., Tumkaya, T., Aryal, S., Choi, H., & Claridge-Chang, A. (2019). Moving beyond P values: data analysis with estimation graphics. Nature Methods, 16, 565–566. https://doi.org/10.1038/s41592-019-0470-3

Kalueff, A. V., Gebhardt, M., Stewart, A. M., Cachat, J. M., Brimmer, M., Chawla, J. S., Craddock, C., Kyzar, E. J., Roth, A., Landsman, S., Gaikwad, S., Robinson, K., Baatrup, E., Tierney, K., Shamchuk, A., Norton, W., Miller, N., Nicolson, T., Braubach, O., … Schneider, and the Z. N. R. C. (ZNRC), Henning. (2013). Towards a Comprehensive Catalog of Zebrafish Behavior 1.0 and Beyond. Zebrafish, 10(1), 70–86. https://doi.org/10.1089/zeb.2012.0861

Lindeyer, C. M., & Reader, S. M. (2010). Social learning of escape routes in zebrafish and the stability of behavioural traditions. Animal Behaviour, 79(4), 827–834. https://doi.org/10.1016/j.anbehav.2009.12.024

Liu, X., Zhang, Y., Lin, J., Xia, Q., Guo, N., & Li, Q. (2016). Social Preference Deficits in Juvenile Zebrafish Induced by Early Chronic Exposure to Sodium Valproate. Frontiers in Behavioral Neuroscience, 10, 201.

Madeira, N., & Oliveira, R. F. (2017). Long-term social recognition memory in zebrafish. Zebrafish, 14, 305–310. http://doi.org/10.1089/zeb.2017.1430

Maruska, K., Soares, M. C., Lima-Maximino, M., Henrique de Siqueira-Silva, D., & Maximino, C. (2019). Social plasticity in the fish brain: Neuroscientific and ethological aspects. Brain Research, 1711, 156–172. https://doi.org/10.1016/j.brainres.2019.01.026

Maximino, C., Oliveira, D. L. de, Rosemberg, D. B., Batista, E. de J. O., Herculano, A. M., Oliveira, K. R. M., Benzecry, R., & Blaser, R. (2012). A comparison of the light/dark and novel tank tests in zebrafish. Behaviour, 149(10–12), 1099–1123. https://doi.org/10.1163/1568539X-00003029

Montgomery, K. C. (1955). The relation between fear induced by novel stimulation and exploratory drive. Journal of Comparative and Physiological Psychology, 48(4), 254–260. https://doi.org/10.1037/h0043788

Montgomery, K. C., & Monkman, J. A. (1955). The relation between fear and exploratory behavior. Journal of Comparative and Physiological Psychology, 48(2), 132–136. https://doi.org/10.1037/h0048596

Norton, W. H. J., Manceau, L., & Reichmann, F. (2019). The Visually Mediated Social Preference Test: A Novel Technique to Measure Social Behavior and Behavioral Disturbances in Zebrafish. Em F. H. Kobeissy (Org.), Psychiatric Disorders: Methods and Protocols (p. 121–132). Springer. https://doi.org/10.1007/978-1-4939-9554-7_8

Nunes, A. R., Carreira, L., Anbalagan, S., Blechman, J., Levkowitz, G., & Oliveira, R. F. (2020). Perceptual mechanisms of social affiliation in zebrafish. Scientific Reports, 10(1), Art. 1. https://doi.org/10.1038/s41598-020-60154-8

Ogi, A., Licitra, R., Naef, V., Marchese, M., Fronte, B., Gazzano, A., & Santorelli, F. M. (2021). Social Preference Tests in Zebrafish: A Systematic Review. Frontiers in Veterinary Science, 7, 590057.

Oliveira, R. (2013). Mind the fish: Zebrafish as a model in cognitive social neuroscience. Frontiers in Neural Circuits, 7, 131.

Parichy, D. M. (2015). Advancing biology through a deeper understanding of zebrafish ecology and evolution. eLife, 4, e05635. https://doi.org/10.7554/eLife.05635

Parra, K. V., Adrian, J. C., & Gerlai, R. (2009). The synthetic substance hypoxanthine 3-N-oxide elicits alarm reactions in zebrafish (Danio rerio). Behavioural Brain Research, 205(2), 336–341. https://doi.org/10.1016/j.bbr.2009.06.037

Pimentel, A. F. N., Lima-Maximino, M. G., Soares, M. C., & Maximino, C. (2021). Zebrafish cooperate while inspecting predators: Experimental evidence for conditional approach. Animal Behaviour, 177, 59–68. https://doi.org/10.1016/j.anbehav.2021.04.014

Ponzoni, L., Sala, M., & Braida, D. (2016). Ritanserin-sensitive receptors modulate the prosocial and the anxiolytic effect of MDMA derivatives, DOB and PMA, in zebrafish. Behavioural Brain Research, 314, 181–189. https://doi.org/10.1016/j.bbr.2016.08.009

Potrich, D., Sovrano, V. A., Stancher, G., & Vallortigara, G. (2015). Quantity discrimination by zebrafish (Danio rerio). Journal of Comparative Psychology, 129(4), 388–393. https://doi.org/10.1037/com0000012

Pritchard, V. L., Lawrence, J., Butlin, R. K., & Krause, J. (2001). Shoal choice in zebrafish, Danio rerio: The influence of shoal size and activity. Animal Behaviour, 62(6), 1085–1088. https://doi.org/10.1006/anbe.2001.1858

Ribeiro, D., Nunes, A. R., Gliksberg, M., Anbalagan, S., Levkowitz, G., & Oliveira, R. F. (2020). Oxytocin receptor signalling modulates novelty recognition but not social preference in zebrafish. Journal of Neuroendocrinology, 32(4), e12834. https://doi.org/10.1111/jne.12834

Richardson, J. T. E. (2011). Eta squared and partial eta squared as measures of effect size in educational research. Educational Research Review, 6(2), 135–147. https://doi.org/10.1016/j.edurev.2010.12.001

Rinkwitz, S., Mourrain, P., & Becker, T. S. (2011). Zebrafish: An integrative system for neurogenomics and neurosciences. Progress in Neurobiology, 93(2), 231–243. https://doi.org/10.1016/j.pneurobio.2010.11.003

Roper, C., & Tanguay, R. L. (2018). Zebrafish as a Model for Developmental Biology and Toxicology. Em W. Slikker, M. G. Paule, & C. Wang (Orgs.), Handbook of Developmental Neurotoxicology (Second Edition) (p. 143–151). Academic Press. https://doi.org/10.1016/B978-0-12-809405-1.00012-2

Rosenthal, G. G., & Ryan, M. J. (2005). Assortative preferences for stripes in danios. Animal Behaviour, 70(5), 1063–1066. https://doi.org/10.1016/j.anbehav.2005.02.005

Saverino, C., & Gerlai, R. (2008). The social zebrafish: Behavioral responses to conspecific, heterospecific, and computer animated fish. Behavioural Brain Research, 191(1), 77–87. https://doi.org/10.1016/j.bbr.2008.03.013

Scerbina, T., Chatterjee, D., & Gerlai, R. (2012). Dopamine receptor antagonism disrupts social preference in zebrafish: A strain comparison study. Amino Acids, 43(5), 2059–2072. https://doi.org/10.1007/s00726-012-1284-0

Seguin, D., & Gerlai, R. (2017). Zebrafish prefer larger to smaller shoals: Analysis of quantity estimation in a genetically tractable model organism. Animal Cognition, 20(5), 813–821. https://doi.org/10.1007/s10071-017-1102-x

Soares, M. C., Cardoso, S. C., Carvalho, T. dos S., & Maximino, C. (2018). Using model fish to study the biological mechanisms of cooperative behaviour: A future for translational research concerning social anxiety disorders? Progress in Neuro-Psychopharmacology and Biological Psychiatry, 82, 205–215. https://doi.org/10.1016/j.pnpbp.2017.11.014

Speedie, N., & Gerlai, R. (2008). Alarm substance induced behavioral responses in zebrafish (Danio rerio). Behavioural Brain Research, 188(1), 168–177. https://doi.org/10.1016/j.bbr.2007.10.031

Stewart, A. M., Braubach, O., Spitsbergen, J., Gerlai, R., & Kalueff, A. V. (2014). Zebrafish models for translational neuroscience research: From tank to bedside. Trends in Neurosciences, 37(5), 264–278. https://doi.org/10.1016/j.tins.2014.02.011

Taborsky, B., & Oliveira, R. F. (2012). Social competence: An evolutionary approach. Trends in Ecology & Evolution, 27(12), 679–688. https://doi.org/10.1016/j.tree.2012.09.003

Taborsky, M., Hofmann, H. A., Beery, A. K., Blumstein, D. T., Hayes, L. D., Lacey, E. A., Martins, E. P., Phelps, S. M., Solomon, N. G., & Rubenstein, D. R. (2015). Taxon matters: Promoting integrative studies of social behavior: NESCent Working Group on Integrative Models of Vertebrate Sociality: Evolution, Mechanisms, and Emergent Properties. Trends in Neurosciences, 38(4), 189–191. https://doi.org/10.1016/j.tins.2015.01.004

